# HIV population-level adaptation can rapidly diminish the impact of a partially effective vaccine

**DOI:** 10.1101/110783

**Authors:** Joshua T. Herbeck, Kathryn Peebles, Paul T. Edlefsen, Morgane Rolland, James T. Murphy, Geoffrey S. Gottlieb, Neil Abernethy, James I. Mullins, John E. Mittler, Steven M. Goodreau

## Abstract

Development of an HIV vaccine is essential to ending the HIV/AIDS pandemic. However, vaccines can result in the emergence and spread of vaccine-resistant strains. Indeed, analyses of breakthrough infections in the HIV vaccine trial RV144 identified HIV genotypes with differential rates of transmission in vaccine and placebo recipients. We hypothesized that, for HIV vaccination programs based on partially effective vaccines similar to RV144, HIV adaptation will diminish the expected vaccine impact. Using two HIV epidemic models, we simulated large-scale vaccination programs and, critically, included HIV strain diversity with respect to the vaccine response. We show here that rapid population-level viral adaptation can lead to decreased overall vaccine efficacy and substantially fewer infections averted by vaccination, when comparing scenarios with and without viral evolution (depending on vaccination coverage, vaccine efficacy against the sensitive allele, and the initial resistant allele frequency). Translating this to the epidemic in South Africa, a scenario with 70% vaccination coverage may result in 250,000 new infections within 10 years of vaccine rollout that are due solely to HIV adaptation, all else being equal. These findings suggest that approaches to HIV vaccine development, program implementation, and epidemic modeling may require attention to viral evolutionary responses to vaccination.

## Main

Despite concerted global effort and the existence of effective methods for prevention, HIV continues to be a public health crisis. The need for an HIV vaccine remains paramount. The phase 3 RV144 HIV vaccine trial is the only trial of an HIV vaccine to show modest success in preventing infection^3^. RV144 resulted in an estimated 31% vaccine efficacy (VE) at 3.5 years post-vaccination (p=0.04, modified intent-to-treat analysis). The vaccine was partially protective but not therapeutic; i.e. vaccinated individuals had decreased rates of infection, but breakthrough infections were not associated with differences in early HIV plasma viral loads, post-infection CD4+ T cell counts, or HIV disease progression rates, when comparing vaccine and placebo recipients^6^. The RV144 results spurred the development of the recently initiated HVTN 702, a large phase 3 HIV vaccine trial in South Africa that aims to replicate the RV144 findings in a different study group, with regimen and schedule that follow from RV144 (with several modifications, including a vaccine insert specific to HIV subtype C, the most common subtype in South Africa).

Partially effective vaccines have been of enduring scientific, clinical, and theoretical interest. For HIV in particular, two decades of mathematical modeling studies suggest that partially effective vaccines, whether protective or therapeutic, can have a substantial impact on the HIV pandemic^7-19^. More recently, HIV epidemic models were used to predict the impact of a partially effective (protective) vaccine similar to RV144 in terms of VE and duration. Models were used to assess the impact of vaccination programs with 30% and 60% population coverage of sexually active adults, with subsequent vaccine rollouts at 1- to 5-year intervals^20^. Results were consistent across several model and epidemic types, e.g., multiple vaccination rounds, at 60% coverage, were predicted to prevent 5-15% of new infections over 10 years^21-28^. The expected impact of vaccination programs depended on vaccination coverage, VE, and the duration of vaccine protection.

However, the potential for HIV adaptation at the population-level in response to vaccination was not considered in these modeling studies. The requirements for adaptive evolution are few: there must be phenotypic variation in a population, this variation must be heritable (linked to genetic variation), and this variation must be related to fitness (differential reproduction)^29^. Evidence from RV144 follow-up studies suggest that, with respect to a partially effective protective vaccine, HIV meets these requirements. Namely, genetic sieve analyses of RV144 breakthrough infections showed that sequences from infected vaccine recipients differed from those isolated from infected placebo recipients. Two signatures were identified in the Env V2 region: in the vaccine recipients, K169X mutations were more frequent (34% vs. 17%) and 181I was more conserved (91% vs. 71%). VE against viruses matching the vaccine at position 169 was 48% (95% confidence interval (CI) 18% to 66%), whereas VE against viruses mismatching the vaccine at position 181 was 78% (CI 35% to 93%)^4,5^. Thus, heritable (genetic) variation in HIV can be associated with differential infection rates in a vaccinated population, making viral adaptation a potential outcome.

We hypothesized that HIV population-level adaptation after vaccine rollout will result from selection acting on a viral locus containing an allele that confers resistance to the vaccine response; i.e., viruses not blocked by a vaccine-elicited immune response will spread in the HIV-infected population. Our goal was to predict the public health impact of this viral evolution, under varying VE, population vaccination coverage and initial frequency of vaccine-resistant genotypes. We quantified this impact in terms of the resistant genotype frequency, the overall VE and the cumulative HIV infections averted by vaccination.

## Methods

We used two stochastic, individual-based HIV epidemic models, both of which were based on existing model frameworks (described in further detail in the Methods, below; Tables S1, S2). The first model was roughly calibrated to the heterosexual epidemic of South Africa^30^, while the second model was parameterized using behavioral data from men who have sex with men (MSM) in the United States. Both models contained antiretroviral therapy (ART) to approximate the population-level effects of ART on background incidence and prevalence in our vaccine simulations, and we repeated all simulations under two ART population coverage levels: 30% and 70% for the heterosexual model, 40% and 70% for the MSM model (lower coverage levels reflect approximate current states; upper level is aspirational).

The vaccine in our models protected individuals from infection by decreasing the per-contact probability of transmission; it did not affect HIV disease progression, viral load, or CD4 count in vaccinated individuals who became infected (consistent with RV144). The mean duration of VE was set to three years with efficacy reduced immediately to 0% at the end of this period. Vaccination programs started in year 20 or 25 (depending on the model), with continuous rollout. Target vaccination coverage was generally reached within three to four years. We modeled viral diversity with respect to vaccine-induced host response via one locus with two alleles: sensitive and resistant. Sensitive viruses had reduced per-contact probabilities of infecting vaccinated hosts. Resistant viruses experienced no change in transmission probability, regardless of the vaccination status of possible recipients. We used a single viral genetic locus as an approximation of the complete set of putative vaccine-resistant variants identified by sieve analyses^4,5^; in effect, multiple independent vaccine-resistant alleles at low frequency will have a similar population-level impact as a single vaccine-resistant allele at high frequency. For each ART coverage level we evaluated two vaccine scenarios, at increasing vaccination coverage levels: 1) VE for the sensitive virus = 75% and initial resistant virus proportion = 0.25; and 2) VE for sensitive virus = 90% and initial resistant virus proportion = 0.50. We selected these scenarios because both correspond to an *overall* VE of roughly ~50% (56.25% and 45%, respectively, calculated as the weighted average of sensitive and resistant VE at first vaccine rollout), a VE that the HVTN 702 vaccine trial is powered to detect relative to the null hypothesis (VE≤25%).

## Results and Discussion

To confirm the validity of our models, we first examined the effects of introducing a partially effective vaccine into HIV epidemics with a single, vaccine-sensitive viral strain (all viruses were equally sensitive to the vaccine response). We found, in agreement with previous models ^21-28^, that a vaccine similar to RV144 can indeed have a modest impact. For example, an HIV vaccine with 45% VE and 50% coverage can prevent ~20% of cumulative infections within 10 years of vaccine rollout, in comparison to populations with no vaccine (Tables S3, S4). As expected, these values increase with higher vaccination coverage and higher VE: a vaccine with 56% VE and 70% overall coverage can prevent ~40% of cumulative infections within 10 years (Tables S3, S4).

Next, we examined the impact of a partially effective vaccine in a population that contained resistant viruses (VE=0 for the resistant viruses). In all simulations, for both heterosexual and MSM models, resistant virus increased in proportion after vaccine rollout (Figure 1). In a representative scenario from the heterosexual model (Figure 1), with 30% ART coverage, 70% vaccination coverage, and 75% VE against the sensitive allele, the resistant allele population proportion increased from 0.25 to 0.38 within 10 years of vaccine rollout. With 70% ART coverage and 90% VE against the sensitive allele, the resistant allele increased from 0.51 to 0.69 within 10 years. Across all epidemic simulations, the rate at which the resistant allele frequency increased was faster with higher vaccination coverage and greater VE against the sensitive virus. As the resistant allele frequency increases, the overall VE correspondingly declines (Figure 2); e.g., with 50% vaccination coverage, overall VE declines from 56% to 50%, or from 45% to 33% in 10 years (depending on sensitive virus VE and initial resistant frequency). Thus, we see that both programmatic and vaccine-related parameters can exert evolutionary pressure on HIV and impact vaccine effectiveness.

**Figure 1.**
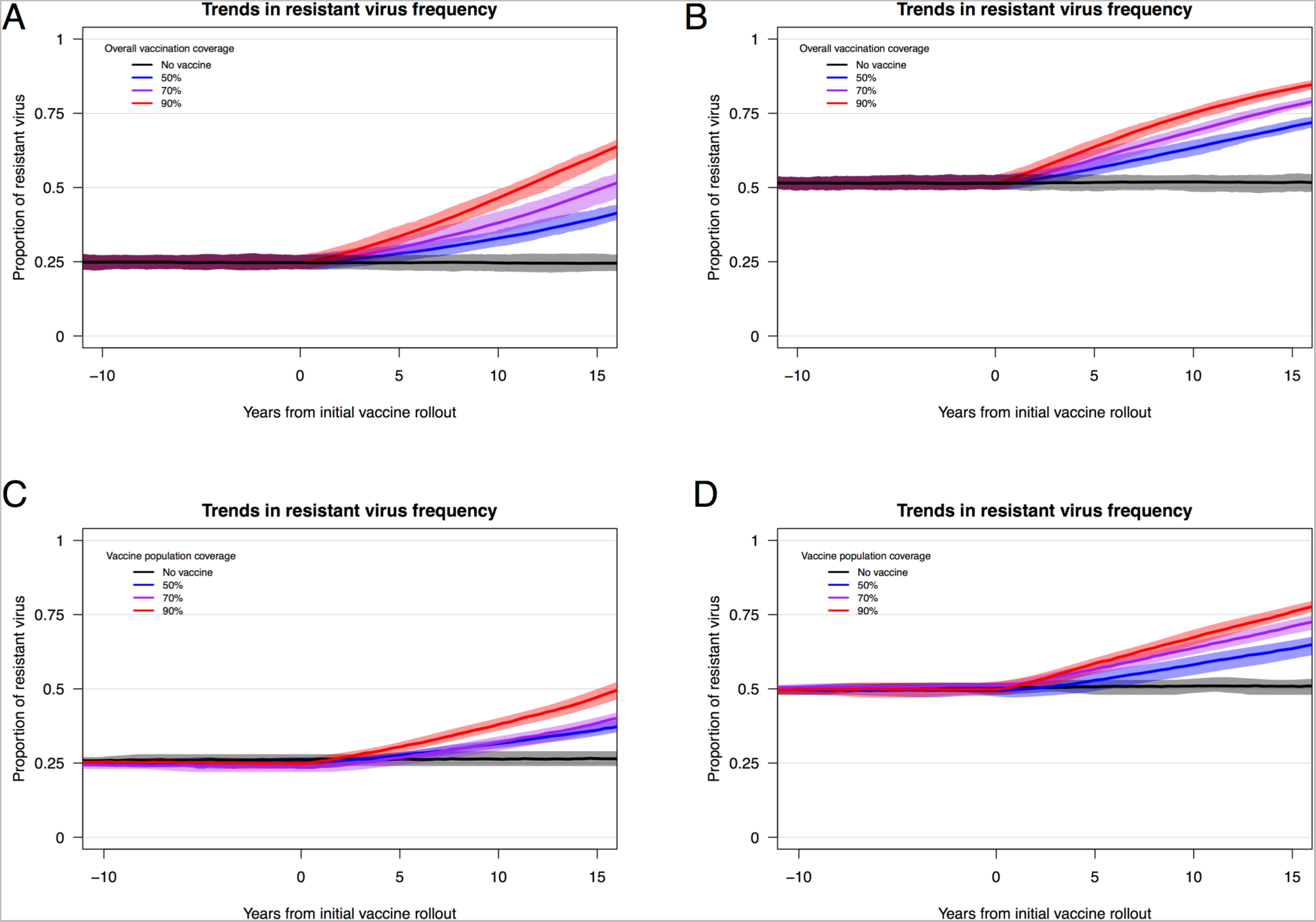
Trends in the frequency of HIV strains that are resistant to a vaccine-driven immune response. Trends in resistant strain proportion (with interquartile ranges) for 20 replicate HIV epidemic simulations from heterosexual (panels A and B) and MSM (panels C and D) models. Panels A and C depict results from epidemic scenarios that included an initial resistant strain (VE = 0.0) proportion = 0.25 and a sensitive virus VE = 0.75. Panels B and D depict results from scenarios with initial resistant strain proportion = 0.50 and a sensitive virus VE = 0.90. Background population ART coverage was at 30% for the heterosexual model and 40% for the MSM model. (See Figure S1 for equivalent results from heterosexual model scenarios in which a high-risk subgroup is preferentially targeted for vaccination, which leads to substantially decreased overall vaccination coverage but results in similar effects of HIV adaptation on resistant virus frequency.)

**Figure 2.**
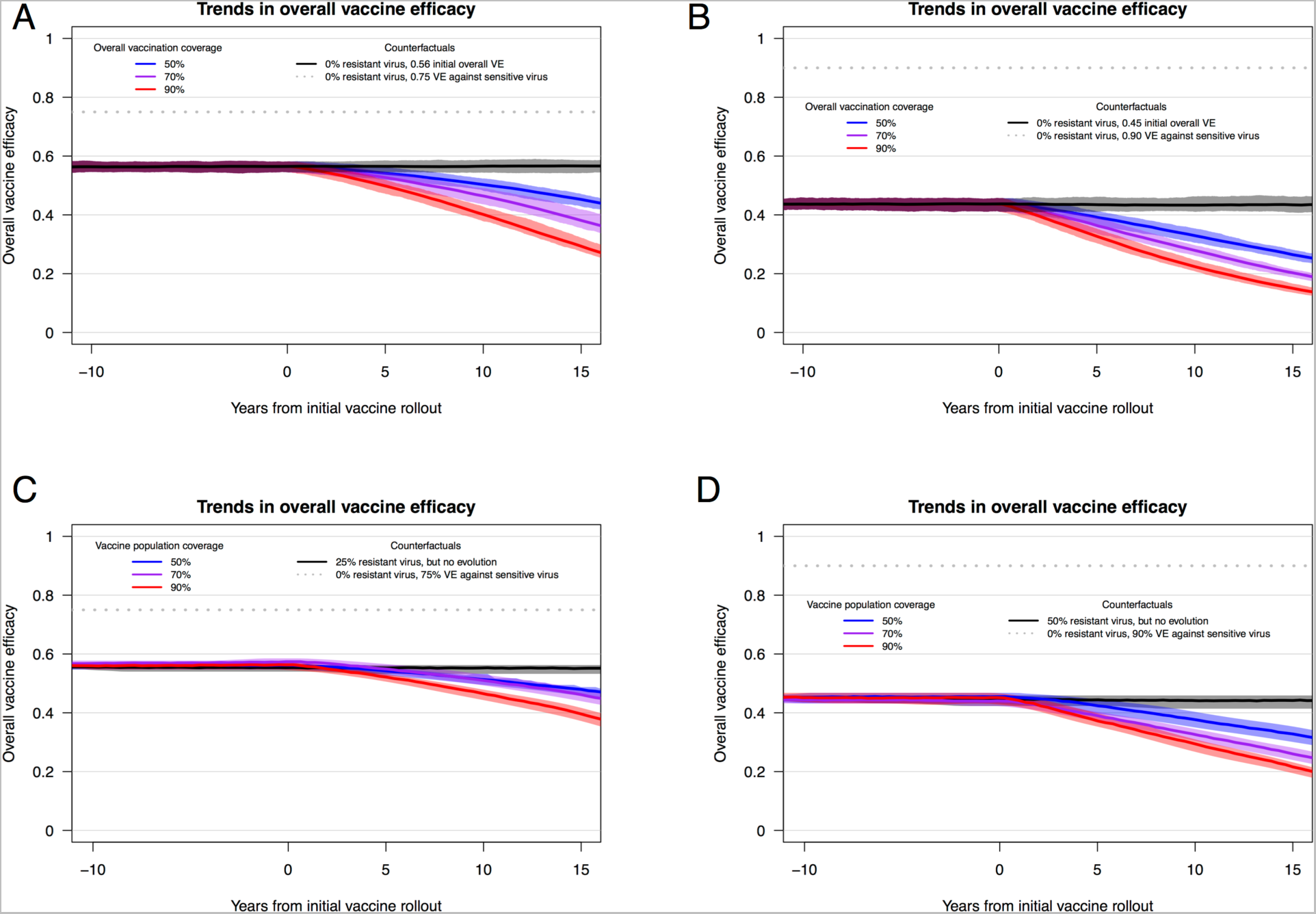
Trends in overall vaccine efficacy (VE) for an HIV vaccine. Trends in overall vaccine efficacy (with interquartile ranges) for 20 replicate HIV epidemic simulations from heterosexual (panels A and B) and MSM (panels C and D) models. Overall VE was calculated as the weighted average of the VE for sensitive and resistant viruses at each time step (VE for resistant viruses = 0.0). Panels A and C depict results from scenarios with an initial resistant strain proportion = 0.25 and a sensitive virus VE = 0.75. Panels B and D depict results from scenarios with initial resistant strain proportion = 0.50 and a sensitive virus VE = 0.90. ART coverage was at 30% for the heterosexual model and 40% for the MSM model. (See Figure S1 and Table S5 for equivalent results from heterosexual model scenarios in which a high-risk subgroup is preferentially targeted for vaccination, which leads to substantially decreased overall vaccination coverage but results in similar effects of HIV adaptation on overall vaccine efficacy.)

### Public health impact

Most critically, our results predict that HIV adaptation in response to vaccination may have a considerable, and detrimental, public health impact. Fewer infections were averted over time in scenarios with resistant strains, relative to counterfactual simulations with no vaccine-resistant HIV strains (Figures 3, 4). Since overall VEs at the time of first vaccine rollout were equivalent across runs, we can assign causality for the marginal differences (in the proportion of infections averted) to viral adaptation. The proportion of infections averted by vaccination decreased dramatically as resistant viruses increased in frequency and overall VE decreased. In the representative examples above, with 70% vaccination coverage and either 75% or 90% VE against the sensitive allele and 25% or 50% resistant virus at vaccine rollout, respectively, infections averted decreased from 39% to 37%, or from 32% to 26%, within 10 years (Figure 3, Table S3). These predictions can be placed in the context of the current HIV epidemic: in South Africa, with approximately 380,000 new HIV infections in 2015^31^, scenarios of 70% vaccination coverage (30% ART coverage, Figure 3) result in approximately 100,000 to 250,000 new infections in the first decade after vaccine rollout that are due solely to the emergence and spread of vaccine-resistant strains (in scenarios of 25% resistant virus and 75% sensitive virus VE, and 50% resistant virus and 90% sensitive virus VE, respectively). These results highlight the potential public health impact of HIV adaptation in response to vaccination. They also underscore the need to understand the underlying viral determinants of partially effective vaccines: despite their similar overall VE of ~50%, the distinct sensitive virus VE and resistant frequency scenarios (25% and 50% resistant virus; 75% and 90% sensitive virus VE) differed greatly in implications for public health.

**Figure 3.**
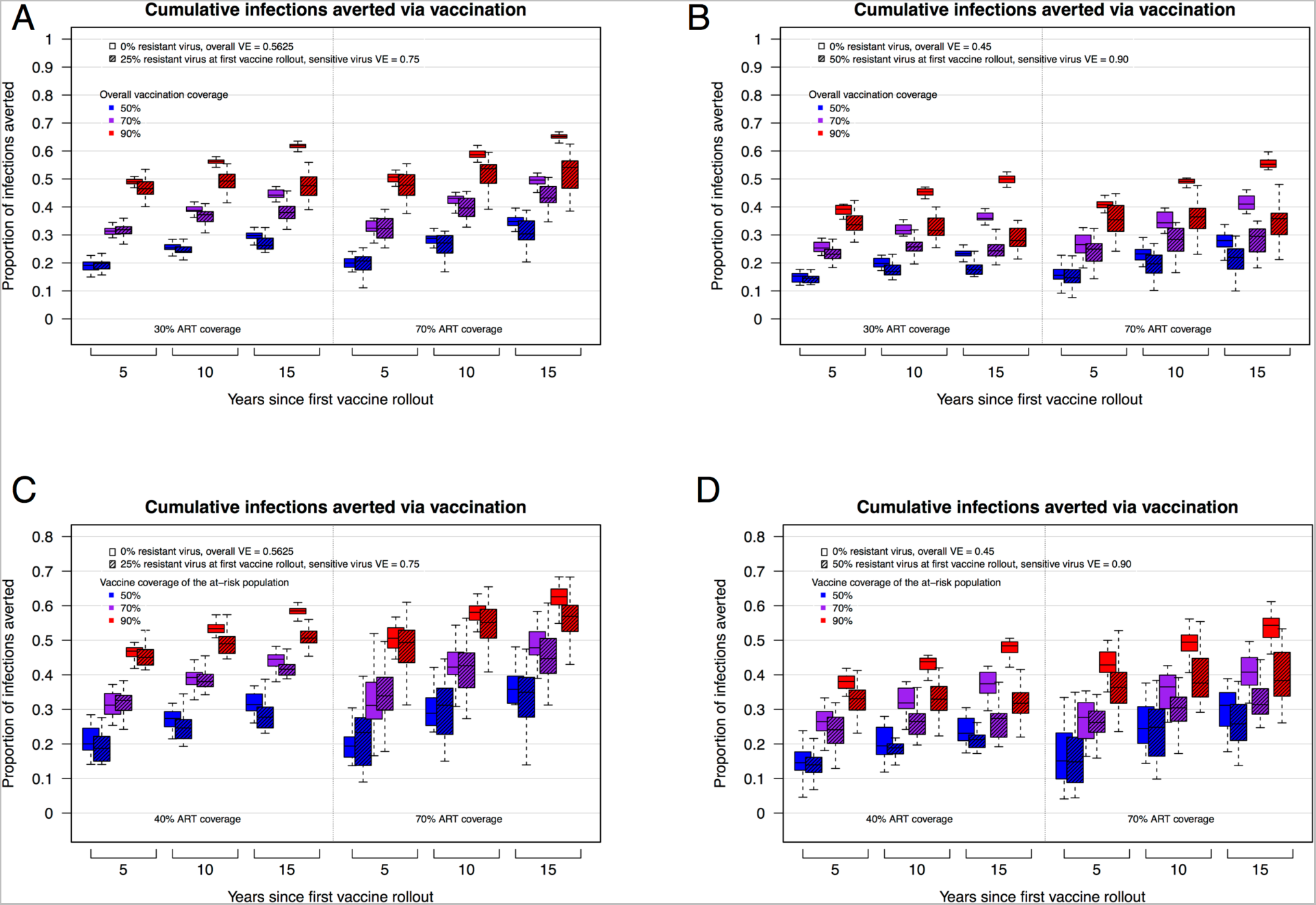
The proportion of cumulative HIV infections averted by vaccination. Trends in cumulative infections averted by vaccination (with interquartile ranges) for 20 replicate epidemic simulations from heterosexual (panels A and B) and MSM (panels C and D) HIV epidemic models (comparing epidemic scenarios with and without vaccination programs). Panels A and C depict results from epidemic scenarios with an initial resistant strain (VE = 0) proportion = 0.25 and a sensitive virus VE = 0.75. Panels B and D depict results with initial resistant strain proportion = 0.50 and a sensitive virus VE = 0.90. All solid color boxplots represent epidemic scenarios with a vaccine but *without* HIV adaptation; the initial overall VE in these scenarios is equal to the overall VE at initial vaccine rollout in the scenarios *with* HIV adaptation (hashed boxplots). ART coverage was at 30% or 70% for the heterosexual model and 40% or 70% for the MSM model. (See Figure S1 and Table S5 for equivalent results from heterosexual model scenarios in which a high-risk subgroup is preferentially targeted for vaccination, which leads to substantially decreased overall vaccination coverage but results in similar effects of HIV adaptation on infections averted.)

In scenarios in which a high-risk subgroup in the heterosexual model is preferentially targeted for vaccination (Figure S1, Table S5), we see equivalent declines in the proportion of infections averted, but with much lower overall vaccination coverage: up to 150,000 new infections in the first 10 years after vaccine rollout due to the emergence and spread of vaccine-resistant strains, in a scenario with 70% vaccination coverage of a high-risk subgroup but only 32% coverage of the total population. This increases to 350,000 new infections with 90% coverage of the high-risk subgroup and 35% total coverage. Vaccination efforts that use high-risk behavioral targeting must be prepared for significant resistance impacts at lower vaccination coverage levels; further studies of the impact of transmission network structure on viral adaptation in response to vaccination, guided by empirical data, are warranted.

We note that similar, but not identical, HIV population-level adaptation and diminished public health impacts were seen in the heterosexual and MSM models, despite differences between the models in population structure, transmission dynamics, HIV prevalence and incidence, and overall effect of vaccination. Additionally, the impacts of HIV adaptation were consistent across the two ART scenarios, suggesting that ART parameters will likely not substantially affect the rates or impact of viral adaptation (Figure 3).

Our HIV-specific model predictions are consistent with findings from other pathogens. These include empirical evidence of vaccine-induced strain replacement in, e.g., *Streptococcus pneumoniae, Haemophilus influenza, Neisseria meningitidis, Bordetella pertussis, Plasmodium falciparum,* and hepatitis B virus (reviewed in^1,2^). Mathematical models have evaluated patterns and processes of strain replacement in, e.g., *Mycobacterium tuberculosis*^32^, rotavirus^33^, and *S. pneumoniae*^34,35^, and generalized pathogens^1,36^ Yet, despite widespread acknowledgement and concern for the evolutionary potential of HIV—with respect to resistance to ART and PrEP^37,38^, vaccine design^39,40^, the human immune response^41-44^, and a potential vaccine-induced cellular immune response^45,46^, HIV strain replacement in response to an imperfect vaccine had not been evaluated previously.

### Conclusions

Previous epidemic models that have estimated the effects of partially effective HIV vaccines have likely overestimated the benefits conferred on a population by vaccination. Perhaps more pressing, strategies for HIV vaccine development and program implementation may benefit from careful attention to the potential evolutionary consequences of vaccination. This includes continued surveillance of viral genetic diversity, accompanied by vaccine design that limits the mutational pathways available for viral adaptation and subsequent emergence of vaccine-resistant viruses^40,47,48^. Our analysis is particularly relevant given the recent initiation of the HVTN 702 trial, which is the critical test for licensure in South Africa of the first vaccine to prevent HIV infection. If successful, such an HIV vaccination program may necessarily evolve into a program similar to that in place for influenza, comprising an acceptable vaccine that requires periodic updating.

## Supplementary Methods (Model descriptions)

Our epidemic models are based on models that have been described previously^49-51^, with modifications to enable simulation of vaccination programs and viral resistance and sensitivity to the vaccine response. These are individual-based stochastic, dynamic models that track inter-host and intra-host dynamics. Agents are endowed with many individual attributes, including demographics, clinical features and behavioral characteristics including sexual role preference (for MSM). HIV-infected individuals possess numerous viral characteristics such as a “set point viral load” (SPVL) and a viral load at each time step over the course of infection. The code from both models is available from the authors upon request (note: all model code will be freely available for review, and will also be deposited on a publically available Github directory, https://github.com/EvoNetHIV, upon publication).

### Heterosexual epidemic model

We simulated a heterosexual HIV epidemic that was calibrated to reproduce incidence and prevalence trajectories based on data from South Africa^52^. We chose to calibrate the model in this way for several reasons: first, because South African has one of the largest HIV epidemics; second, because the large-scale HIV vaccine trial HVTN 702 is located in South Africa, and is based upon the RV144 trial design and results; and third, so that the epidemic output was directly comparable to the 12 HIV epidemic models described in Eaton et al.^30^ Using an approach common among those models, we first set empirical parameter values (i.e., viral load trajectories, CD4 progression rates, set point viral load distribution), then subsequently calibrated the assumption-based parameter values to fit expected epidemic dynamics. Parameters and initial values are listed in the Table S1.

For each individual, upon HIV-1 infection with an initial viral population size, viral load increased exponentially until a peak viremia at the midpoint of the length of acute infection. Viral load then declined exponentially at an individual-specific decay rate until it reached set point viral load (SPVL). After reaching set point, the viral load increased in a log linear manner until the onset of AIDS. Viral load upon onset of AIDS is defined as the same for all individuals, and is independent of SPVL. To model the relationship between SPVL and disease progression, the model related individual SPVL to the starting CD4 count category (individual CD4 count immediately after infection) and to disease progression based on waiting times in four CD4 count categories: CD4≥500; 500>CD4≥350; 350>CD4≥200; CD4<200 (AIDS)^53^.

Transmission rates were assumed to follow available data from serodiscordant heterosexual partners; the probability that an HIV-infected person will transmit to an HIV-negative person is determined by a Hill function that follows from Fraser^54^. The individual SPVL is determined by both viral and environmental factors, with the viral component being heritable across transmissions. For each individual in a simulation, the model maintains a list of sexual partnerships and viral transmission pairs.

For any one epidemic simulation, an initial set of contacts was formed by randomly choosing pairs from the population, prior to the simulation of viral transmission. The probability of each person entering this link was set to *e*^-Li^, where *L*_i_ is the number contacts for person *i*. If this probability was not met (or if *L*_i_ > *MaxLinks*), another partner was selected at random from the population. This process was repeated until the total number of links equals *N*\**M*/2, where *N* is the population and *M* is the mean degree. The probability of a connection between individuals *i* and *j* dissolving is set to 2/(*Duration*[*i*]* *Duration*[*j*]), where *Duration*[*x*] is the expected time that person *x* stays in a relationship.

The viral and CD4-based progression functions are embedded in a host population without demographic (sex, age) heterogeneity, but with behavioral heterogeneity, using a framework adapted from previous work. Each individual is assigned a relational duration propensity; when two individuals partner, their relationship lasts for the mean of these individual effects. The daily probability of sex according to relationship duration; from this variation, we established three subpopulations with distinct risk profiles defined by relationship duration and daily probability of sexual contact. The high-risk group was defined by the shortest mean relationship duration (<0.5 years) and the highest daily probability of sex (0.8); the frequency of this high-risk subgroup was 15% (15% of the initial population, and 15% of all new individual entries into the population). Daily probability of sex for the two lower-risk subgroups were 0.12 and 0.08 (6.67% and 10% reductions in sexual activity, as a proxy for transmission and infection risk, compared to the high-risk subgroup). We did not include a change in behavior patterns over calendar time; e.g., we did not include a reduction in the average sexual contact rate, across individuals, as the epidemic progresses.

We ran the heterosexual epidemic simulation for 40 years, to mimic HIV epidemics from 1990 to 2030 in South Africa. We calibrated epidemic simulations to begin with 2% HIV incidence (per 100 person years), which rose quickly to ~3%, followed by a decline to ~1.5% and stabilization by 2010. Prevalence rose to ~13%, followed by a decline to ~10% with ART coverage at approximately 30% (Figure 4).

**Figure 4.**
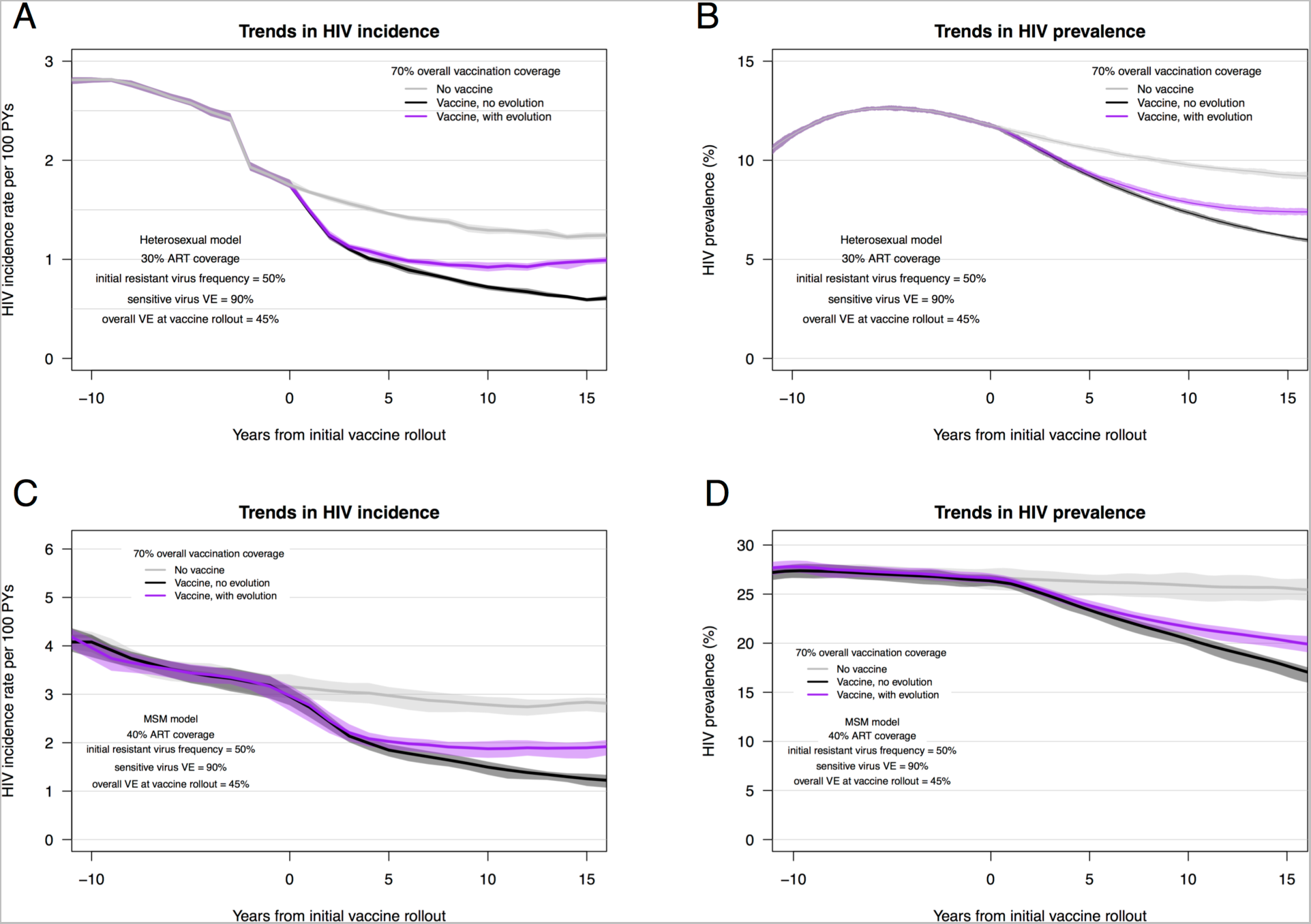
Trends in the incidence and prevalence for the HET and MSM HIV epidemic models, with and without HIV vaccination. Trends in HIV incidence (with interquartile ranges) for 20 replicate HIV epidemic simulations from heterosexual (panel A) and MSM (panel C) models. Trends in HIV prevalence from heterosexual (panel B) and MSM (panel D) models. All panels depict results from epidemic scenarios that included an initial resistant strain (VE = 0) proportion = 0.50 and a sensitive virus VE = 0.90. Background population ART coverage was at 30% for the heterosexual model and 40% for the MSM model. (See Figure S1 and Table S5 for equivalent results from heterosexual model scenarios in which a high-risk subgroup is preferentially targeted for vaccination, which leads to substantially decreased overall vaccination coverage but results in similar effects of HIV adaptation on incidence and prevalence.)

### Men-who-have-sex-with-men (MSM) epidemic model

In the MSM HIV epidemic model, the dynamics of sexual relationships were governed by network models based in the separable-temporal exponential-family random graph model (STERGM) framework^55,56^.

Each simulation started with a population of HIV-uninfected and infected individuals at time zero. Dyads formed and dissolved partnerships to match target network parameters. Within ongoing partnerships, we modeled individual coital acts, as well as decisions about condom use per act. For each serodiscordant coital act that occurred, the probability of transmission from an HIV-infected person to an HIV-negative person was based primarily on viral load but also on disease stage (above and beyond viral load), condom use, and antiretroviral use; this is described by a complementary-log log link function that follows from Hughes et al.^57^.

Individual viral load trajectories were modeled with five phases in the absence of treatment: an initial increase from infection to acute peak; a two-phase decay to set point viral load; a long, chronic phase with slow viral load increase, and a final AIDS stage. Individuals varied in their set point viral load, with individual values set to have both a heritable component from their infector, and a random (environmental) component.

The MSM model varied in several key ways from the source model^51,55^, as befits the aims of this work. We simplified from three separate network models (for main, casual, and one-time partners) in those models to just one overall network per model, since we were not focused on understanding the evolutionary response to vaccination stratified by partner and exposure types. Four features already mentioned—variation in SPVL, post-acute biphasic decay in viral load, administration of vaccine, and the existence and heritability of vaccine resistance—were new to this model. Finally, because of these changes the model contains numerous novel parameters that were derived from the data described in Table S2.

The complete code (called EvoNetHIV) is built as a series of R function modules that utilize the epidemic modeling framework EpiModel^58^, itself an R package. An additional detailed technical supplement is available at https://github.com/EvoNetHIV.

### Vaccine programs

The vaccine parameters related to duration and the effect on transmission/infection are identical in both of our models, and are as described in the main text. 20 replicate simulations were performed for each combination of parameters for both models. For the MSM simulations, the vaccination program was initiated 10 years after equilibrium prevalence of ~25% was reached; this was generally around year 20 of the simulations. For the heterosexual simulations, the vaccination program was initiated at year 25 of the simulated epidemic. Vaccine rollout was continuous after initiation; target coverage was generally reached within three to four years. Across viral transmissions there was 100% heritability of the vaccine-related genotype, and no within-host viral evolution took place that affected the genotype. Individuals who are already infected were not eligible for vaccination.

As described in the main text, the duration of vaccine efficacy (VE) was three years, with VE reduced immediately to 0% at the end of this period; while VE in RV144 declined from ~60% to ~30% over 3.5 years, we chose to employ a simplified flat VE trajectory, as recent RV144-related modeling work by Gilbert et al. has shown that the shape of the temporal VE profile has a negligible, and statistically insignificant, impact on model results^59^.

For the heterosexual model, we employed two separate approaches to vaccination: random and targeted. For random vaccination, individuals had the same probability of being vaccinated, regardless of their risk group status. In this case, population vaccination coverage was a single target and reflected the entire population of susceptible individuals. For targeted vaccination, the high-risk subgroup was given a significantly higher target vaccination coverage (50%, 70%, 90%), while the two lower-risk subgroups were both given 25% vaccination coverage targets.

### Antiretroviral therapy

Antiretroviral therapy (ART) became available in 2012 in the heterosexual South African model, and was available at all time points in the MSM model. In both models, individuals became eligible for treatment after they had been infected for a mean of three years, regardless of their CD4 count. In the MSM model, this parameter was set to a mean of three years; in the HET model this time until eligibility for ART was set to exactly three years for all individuals, with no distribution around this time. This time-since-infection eligibility criterion is meant to approximate “test and treat” guidelines, but scaled to reflect delayed diagnosed and treatment initiation, in both the USA and sub-Saharan Africa. Population coverage was set at approximately 30% and 40%, for the HET and MSM models, respectively, and 70% for both models, for comparison. The 70% high coverage, with assumed complete adherence and viral suppression, is meant to approximate the UNAIDS 90-90-90 goals, which if implemented successfully would lead to 72% of all infected individuals receiving therapy and becoming virally suppressed.

We repeated all vaccine scenarios for both ART coverage levels scenarios; all ART scenarios were represented in the counterfactual epidemic scenarios (no vaccine and vaccine with no viral variation) and the epidemic scenarios that included viral resistance to the vaccine effect.

## Supplementary Materials

File includes Tables S1 – S5 and Figure S1

## Acknowledgments

We thank Jonathan Carlson, Christophe Fraser, Christian Selinger and members of the University of Washington International Clinical Research Center for input and discussion. This work was supported by grants from the U.S. National Institutes of Health (R01AI108490 to J.T.H., J.E.M., and S.M.G., and P30AI027757 to the University of Washington Center for AIDS Research) and by an Interagency Agreement with the US Army Medical Research and Material Command (Y1-AI-2642-12) and by a cooperative agreement between the Henry M. Jackson Foundation for the Advancement of Military Medicine, Inc., and the US Department of Defense (MR). The content is solely the responsibility of the authors and does not necessarily represent the official views of the National Institutes of Health, the US Department of Defense or the Department of the Army. The authors declare no competing interests.

## Author contributions

J.T.H. conceived the study and designed the experiments. J.T.H., K.P., J.T.M., G.S.G., N.A., J.E.M, and S.G.M. developed the modeling platforms. J.T.H and K.P. performed the modeling experiments. J.T.H., K.P., P.T.E., M.R., G.S.G., N.A., J.I.M, J.E.M, and S.M.G. interpreted the data and contributed to writing the manuscript.

---

**Table S1.** Parameters of the heterosexual HIV epidemic model and initial values.

| Parameter | Value |
| --- | --- |
| **Demographic and behavioral** |  |
| Initial overall population size | $1 \times 10^5$ individuals |
| Initial number of infected | $2 \times 10^3$ individuals |
| Minimum relationship duration | 0.1 years |
| Maximum relationship duration | 5.0 years |
| Subgroups defined by relationship duration | <0.5 yrs, 0.5-2.5 yrs, >2.5 yrs |
| Probability of sexual contact, in each group | 0.8, 0.12, 0.08 per day |
| Mean degree | 0.9 |
| **Virologic** |  |
| Viral load at time zero | 10 copies/mL |
| Viral load at peak viremia | $1.0 \times 10^7$ copies/mL<sup>60,61</sup> |
| Time to peak viremia | 21 days<sup>60,61</sup> |
| Total time of acute infection | 91 days<sup>60</sup> |
| Viral load progression rate, natural log | 0.14 per year<sup>62</sup> |
| Viral load at AIDS (CD4<200) | $5.0 \times 10^6$ copies/mL<sup>62</sup> |
| **Set point viral load** |  |
| Variance of $\log_{10}$ SPVL | 0.7<sup>63,64</sup> |
| Heritability of SPVL across transmissions ($h^2$) | 0.36<sup>65</sup> |
| **Transmission** |  |
| Maximum transmission rate | 0.0055 per day<sup>54</sup> |
| Viral load at 0.5 max transmission rate | 13,938 copies/mL<sup>54</sup> |
| Hill coefficient, transmission function | 1.02<sup>54</sup> |
| Shape parameter, transmission function | 3.46<sup>54</sup> |
| **Disease progression** |  |
| See Cori *et al.* for CD4 wait time matrix stratified by SPVL<sup>53</sup> |  |
| **Antiretroviral therapy** |  |
| Time to initiation after becoming eligible by CD4 | 1 year<sup>30</sup> |
| Viral load after ART initiation | 50 copies/mL |
| **Vaccine** |  |
| Vaccine efficacy (of the sensitive allele) | 0.50, 0.75 |
| Duration of vaccine efficacy | 3 years<sup>3</sup> |

**Table S2.** Parameters of the men-who-have-sex-with-men HIV epidemic model and initial values.

| Parameter | Value |
| --- | --- |
| **Demographic and behavioral** |  |
| Initial overall population size | 10,000 |
| Initial number of infected | 2,000 |
| Age range | 18-55 |
| Mean relationship duration | 50 days |
| Mean degree | 0.7 |
| Daily probability of sexual contact (in a relationship) | 0.2<sup>51</sup> |
| Condom use probability per sexual contact | 0.5<sup>51</sup> |
| Proportion of population circumcised | 0.85<sup>51</sup> |
| **Virologic** |  |
| Viral load at day 1 of infection | 1.0x10<sup>-4</sup> |
| Viral load at peak viremia | 7.7x10<sup>6</sup> copies/mL<sup>66</sup> |
| Time to peak viremia | 21 days<sup>60,61</sup> |
| Total time of acute infection | 90 days<sup>67</sup> |
| Viral load progression rate, natural log | 0.14 per year<sup>62</sup> |
| Maximum viral load in AIDS (CD4<200) | 2.4x10<sup>6</sup> copies/mL<sup>68</sup> |
| **Set point viral load** |  |
| Variance of log<sub>10</sub> SPVL | 0.8<sup>63</sup> |
| Heritability of SPVL across transmissions (h<sup>2</sup>) | 0.36<sup>65</sup> |
| Mutational variance | 0.01 |
| **Transmission** |  |
| Relative risk per log<sub>10</sub> increase in viral load | 2.89<sup>57</sup> |
| Relative risk of insertive anal intercourse | 2.9<sup>69</sup> |
| Relative risk of receptive anal intercourse | 17.3<sup>69</sup> |
| Relative risk of condom use | 0.22<sup>57</sup> |
| Relative risk of circumcision | 0.53<sup>57</sup> |
| **Disease progression** |  |
| See Cori *et al.* for CD4 wait time matrix stratified by SPVL<sup>53</sup> |  |
| **Antiretroviral therapy** |  |
| ART coverage of eligible individuals | 40% |
| CD4 threshold for treatment eligibility | <350 cells/mm<sup>3</sup> |
| HIV testing interval | 365 days<sup>70</sup> |
| Viral load after ART initiation | 13 copies/mL<sup>71</sup> |

**Table S3.** Proportion of cumulative HIV infections averted by vaccination. Proportion of cumulative infections averted, for the heterosexual model with 30% ART coverage (median (minimum - maximum)) and two scenarios of initial frequency of resistant virus (25% or 50%) and vaccine efficacy against the sensitive virus (75% and 90%).

| Overall vaccination coverage |  |  |  |
| --- | --- | --- | --- |
|  | 50% | 70% | 90% |
| **No resistant viral genotypes (overall VE = 0.5625)** |  |  |  |
| 5 years | 0.19 (0.15 - 0.23) | 0.31 (0.29 - 0.35) | 0.49 (0.47 - 0.51) |
| 10 years | 0.25 (0.22 - 0.28) | 0.39 (0.36 - 0.42) | 0.56 (0.54 - 0.59) |
| 15 years | 0.29 (0.26 - 0.33) | 0.44 (0.42 - 0.47) | 0.62 (0.60 - 0.64) |
| **25% resistant viruses at start (overall VE at vaccine rollout = 0.5625)\*** |  |  |  |
| 5 years | 0.20 (0.16 - 0.23) | 0.32 (0.27 - 0.36) | 0.47 (0.40 - 0.54) |
| 10 years | 0.24 (0.21 - 0.30) | 0.37 (0.31 - 0.41) | 0.49 (0.42 - 0.55) |
| 15 years | 0.26 (0.24 - 0.33) | 0.38 (0.32 - 0.46) | 0.47 (0.39 - 0.56) |
| **\*Vaccine efficacy for sensitive allele = 75%** |  |  |  |
| Overall vaccination coverage |  |  |  |
|  | 50% | 70% | 90% |
| **No resistant viral genotypes (overall VE = 0.45)** |  |  |  |
| 5 years | 0.15 (0.12 - 0.18) | 0.25 (0.23 - 0.29) | 0.39 (0.36 - 0.41) |
| 10 years | 0.20 (0.17 - 0.23) | 0.32 (0.30 - 0.36) | 0.45 (0.43 - 0.47) |
| 15 years | 0.23 (0.20 - 0.26) | 0.36 (0.34 - 0.39) | 0.50 (0.47 - 0.53) |
| **50% resistant viruses at start (overall VE at vaccine rollout = 0.45)\*** |  |  |  |
| 5 years | 0.15 (0.09 - 0.21) | 0.23 (0.18 - 0.29) | 0.34 (0.27 - 0.42) |
| 10 years | 0.17 (0.14 - 0.23) | 0.26 (0.20 - 0.32) | 0.32 (0.26 - 0.40) |
| 15 years | 0.18 (0.15 - 0.24) | 0.24 (0.19 - 0.32) | 0.28 (0.21 - 0.35) |
| **\*Vaccine efficacy for sensitive allele = 90%** |  |  |  |

**Table S4.** Proportion of cumulative HIV infections averted by vaccination. Proportion of cumulative infections averted, for the MSM model with 40% ART coverage (median (minimum - maximum)) and two scenarios of initial frequency of resistant virus (25% or 50%) and vaccine efficacy against the sensitive virus (75% and 90%).

| Overall vaccination coverage |  |  |  |
| --- | --- | --- | --- |
|  | 50% | 70% | 90% |
| **No resistant viral genotypes (overall VE = 0.5625)** |  |  |  |
| 5 years | 0.20 (0.14 - 0.28) | 0.31 (0.25 - 0.37) | 0.47 (0.41 - 0.53) |
| 10 years | 0.27 (0.21 - 0.32) | 0.39 (0.33 - 0.44) | 0.53 (0.49 - 0.57) |
| 15 years | 0.31 (0.26 - 0.37) | 0.45 (0.38 - 0.48) | 0.58 (0.55 - 0.63) |
| **25% resistant viruses at start (overall VE at vaccine rollout = 0.5625)\*** |  |  |  |
| 5 years | 0.19 (0.14 - 0.28) | 0.33 (0.24 - 0.38) | 0.45 (0.41 - 0.53) |
| 10 years | 0.25 (0.19 - 0.36) | 0.38 (0.29 - 0.46) | 0.49 (0.45 - 0.57) |
| 15 years | 0.28 (0.23 - 0.40) | 0.42 (0.32 - 0.48) | 0.51 (0.44 - 0.56) |
| **\*Vaccine efficacy for sensitive allele = 75%** |  |  |  |
| Overall vaccination coverage |  |  |  |
|  | 50% | 70% | 90% |
| **No resistant viral genotypes (overall VE = 0.45)** |  |  |  |
| 5 years | 0.15 (0.05 - 0.27) | 0.26 (0.18 - 0.33) | 0.38 (0.30 - 0.48) |
| 10 years | 0.19 (0.12 - 0.28) | 0.32 (0.24 - 0.38) | 0.44 (0.36 - 0.52) |
| 15 years | 0.23 (0.17 - 0.30) | 0.37 (0.30 - 0.42) | 0.48 (0.42 - 0.57) |
| **50% resistant viruses at start (overall VE at vaccine rollout = 0.45)\*** |  |  |  |
| 5 years | 0.14 (0.07 - 0.28) | 0.24 (0.13 - 0.32) | 0.33 (0.23 - 0.46) |
| 10 years | 0.19 (0.14 - 0.30) | 0.27 (0.20 - 0.36) | 0.33 (0.22 - 0.42) |
| 15 years | 0.21 (0.17 - 0.29) | 0.27 (0.19 - 0.38) | 0.32 (0.22 - 0.42) |
| **\*Vaccine efficacy for sensitive allele = 90%** |  |  |  |

**Table S5.** Proportion of cumulative HIV infections averted by targeted vaccination. Targeted vaccination of a high-risk subgroup, for the heterosexual model with 30% ART coverage (median (minimum - maximum)) and two scenarios of initial frequency of resistant virus (25% or 50%) and vaccine efficacy against the sensitive virus (75% and 90%).

| Vaccination coverage (high-risk subgroup; overall pop.) |  |  |  |
| --- | --- | --- | --- |
|  | 50%; 29% | 70%; 32% | 90%; 35% |
| **No resistant viral genotypes (overall VE = 0.5625)** |  |  |  |
| 5 years | 0.14 (0.10 – 0.17) | 0.20 (0.16 – 0.24) | 0.29 (0.26 – 0.44) |
| 10 years | 0.19 (0.15 – 0.22) | 0.28 (0.24 – 0.30) | 0.37 (0.34 – 0.39) |
| 15 years | 0.23 (0.18 – 0.26) | 0.33 (0.30 – 0.35) | 0.44 (0.41 – 0.46) |
| **25% resistant viruses at start (overall VE at vaccine rollout = 0.5625)\*** |  |  |  |
| 5 years | 0.15 (0.08 – 0.18) | 0.20 (0.15 – 0.24) | 0.28 (0.23 – 0.32) |
| 10 years | 0.19 (0.13 – 0.23) | 0.25 (0.20 – 0.29) | 0.32 (0.27 – 0.37) |
| 15 years | 0.21 (0.17 – 0.27) | 0.28 (0.23 – 0.32) | 0.33 (0.25 – 0.38) |
| **\*Vaccine efficacy for sensitive allele = 75%** |  |  |  |
| Vaccination coverage (high-risk subgroup; overall pop.) |  |  |  |
|  | 50%; 29% | 70%; 32% | 90%; 35% |
| **No resistant viral genotypes (overall VE = 0.45)** |  |  |  |
| 5 years | 0.11 (0.08 – 0.15) | 0.17 (0.14 – 0.19) | 0.23 (0.21 – 0.26) |
| 10 years | 0.16 (0.14 – 0.19) | 0.22 (0.19 – 0.24) | 0.30 (0.27 – 0.31) |
| 15 years | 0.18 (0.16 – 0.21) | 0.26 (0.24 – 0.29) | 0.35 (0.33 – 0.38) |
| **50% resistant viruses at start (overall VE at vaccine rollout = 0.45)\*** |  |  |  |
| 5 years | 0.12 (0.07 – 0.15) | 0.14 (0.12 – 0.19) | 0.20 (0.17 – 0.26) |
| 10 years | 0.14 (0.10 – 0.17) | 0.18 (0.13 – 0.23) | 0.21 (0.18 – 0.27) |
| 15 years | 0.14 (0.11 – 0.19) | 0.18 (0.12 – 0.23) | 0.19 (0.16 – 0.25) |
| **\*Vaccine efficacy for sensitive allele = 90%** |  |  |  |

**Figure S1.**
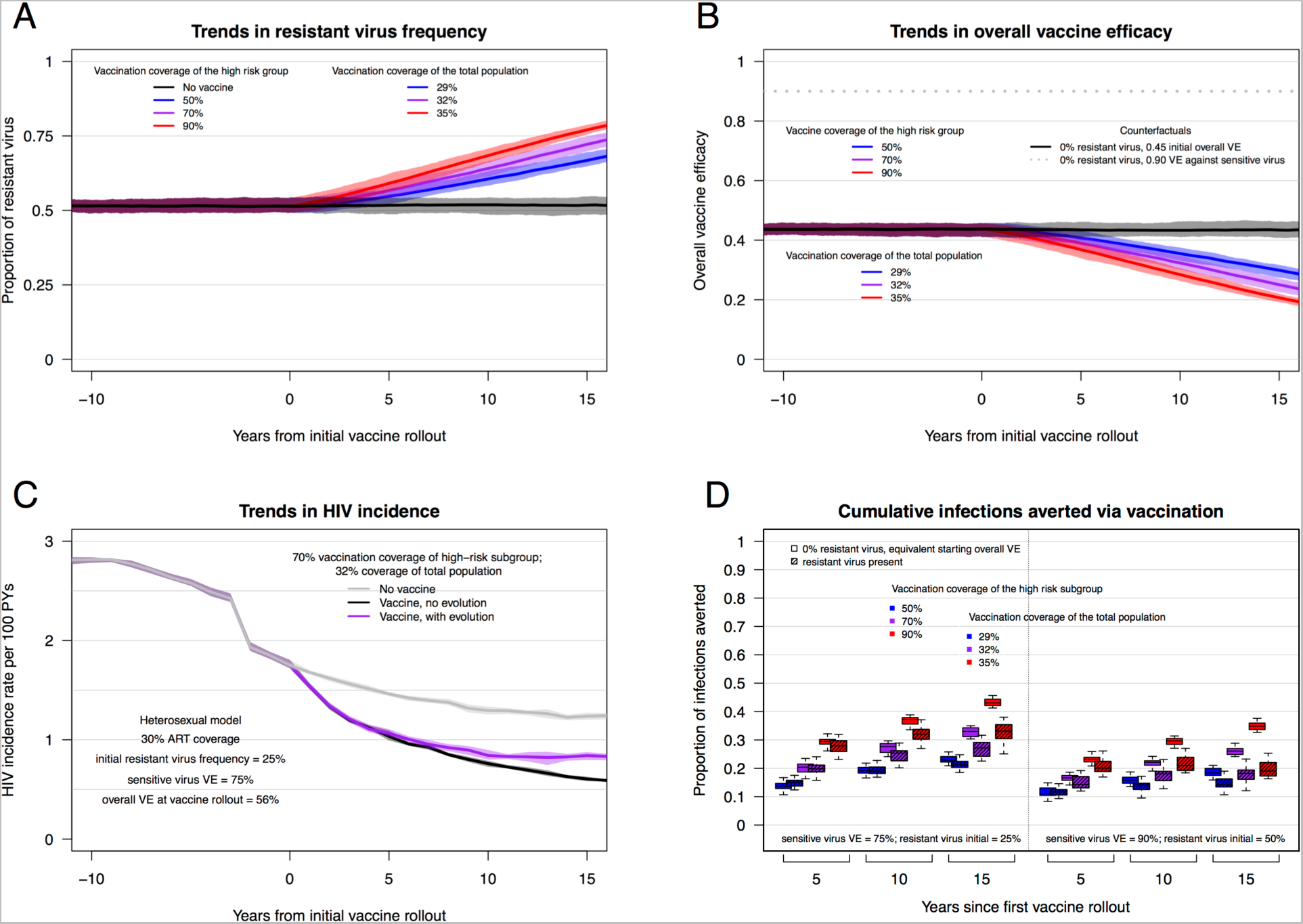
Effects of targeted vaccination on a high-risk subgroup of the total population. Trends in resistant allele frequency, overall vaccine efficacy, HIV incidence, and cumulative infections averted, for epidemic scenarios in which a high-risk subgroup (representing ~15% of the population) is preferentially targeted for vaccination. Vaccination coverage of the high-risk subgroup is relatively high (>50%), but overall vaccination coverage is low (<50%); effects of HIV adaptation in response to vaccination are similar to results (presented above) that include relatively high overall vaccination coverage. All panels depict results from epidemic scenarios with an initial resistant strain (VE = 0) proportion = 0.50 and a sensitive virus VE = 0.90; panel D shows results from this scenario and also the scenario with initial resistant strain (VE = 0) proportion = 0.25 and a sensitive virus VE = 0.75

## References

1. Martcheva, M., Bolker, B.M. & Holt, R.D. Vaccine-induced pathogen strain replacement: what are the mechanisms? J R Soc Interface 5, 3–13 (2008).

2. Read, A.F. & McKinnon, M.J. in Evolution in Health and Disease (eds. Stearns, S.C. & Koella, J.) 139–152 (Oxford University Press, 2008).

3. Rerks-Ngarm, S., et al. Vaccination with ALVAC and AIDSVAX to prevent HIV-1 infection in Thailand. The New England journal of medicine 361, 2209–2220 (2009).

4. Edlefsen, P.T., et al. Comprehensive sieve analysis of breakthrough HIV-1 sequences in the RV144 vaccine efficacy trial. PLoS Comput Biol 11, e1003973 (2015).

5. Rolland, M., et al. Increased HIV-1 vaccine efficacy against viruses with genetic signatures in Env V2. Nature 490, 417–420 (2012).

6. Rerks-Ngarm, S., et al. Extended evaluation of the virologic, immunologic, and clinical course of volunteers who acquired HIV-1 infection in a phase III vaccine trial of ALVAC-HIV and AIDSVAX B/E. J Infect Dis 207, 1195–1205 (2013).

7. Andersson, K.M., et al. Predicting the impact of a partially effective HIV vaccine and subsequent risk behavior change on the heterosexual HIV epidemic in low-and middle-income countries: A South African example. Journal of acquired immune deficiency syndromes (1999) 46, 78–90 (2007).

8. Long, E.F., Brandeau, M.L. & Owens, D.K. Potential population health outcomes and expenditures of HIV vaccination strategies in the United States. Vaccine 27, 5402–5410 (2009).

9. Davenport, M.P., Ribeiro, R.M., Chao, D.L. & Perelson, A.S. Predicting the impact of a nonsterilizing vaccine against human immunodeficiency virus. J Virol 78, 11340–11351 (2004).

10. Owens, D.K., Edwards, D.M. & Shachter, R.D. Population effects of preventive and therapeutic HIV vaccines in early- and late-stage epidemics. AIDS 12, 1057–1066 (1998).

11. Abu-Raddad, L.J., Boily, M.C., Self, S. & Longini, I.M. Jr., Analytic insights into the population level impact of imperfect prophylactic HIV vaccines. Journal of acquired immune deficiency syndromes (1999) 45, 454–467 (2007).

12. Smith, R.J. & Blower, S.M. Could disease-modifying HIV vaccines cause population-level perversity? The Lancet. Infectious diseases 4, 636–639 (2004).

13. Anderson, R. & Hanson, M. Potential public health impact of imperfect HIV type 1 vaccines. J Infect Dis 191 Suppl 1, S85–96 (2005).

14. van Ballegooijen, M., Bogaards, J.A., Weverling, G.J., Boerlijst, M.C. & Goudsmit, J. AIDS vaccines that allow HIV-1 to infect and escape immunologic control: a mathematic analysis of mass vaccination. Journal of acquired immune deficiency syndromes (1999) 34, 214–220 (2003).

15. Anderson, R.M., Swinton, J. & Garnett, G.P. Potential impact of low efficacy HIV-1 vaccines in populations with high rates of infection. Proc Biol Sci 261, 147–151 (1995).

16. McLean, A.R. & Blower, S.M. Imperfect vaccines and herd immunity to HIV. Proc Biol Sci 253, 9–13 (1993).

17. Amirfar, S., Hollenberg, J.P. & Abdool Karim, S.S. Modeling the impact of a partially effective HIV vaccine on HIV infection and death among women and infants in South Africa. Journal of acquired immune deficiency syndromes (1999) 43, 219–225 (2006).

18. Gray, R.H., et al. Stochastic simulation of the impact of antiretroviral therapy and HIV vaccines on HIV transmission; Rakai, Uganda. AIDS 17, 1941–1951 (2003).

19. Blower, S.M. & McLean, A.R. Prophylactic vaccines, risk behavior change, and the probability of eradicating HIV in San Francisco. Science 265, 1451–1454 (1994).

20. Hankins, C.A., Glasser, J.W. & Chen, R.T. Modeling the impact of RV144-like vaccines on HIV transmission. Vaccine 29, 6069–6071 (2011).

21. Andersson, K.M., Paltiel, A.D. & Owens, D.K. The potential impact of an HIV vaccine with rapidly waning protection on the epidemic in Southern Africa: examining the RV144 trial results. Vaccine 29, 6107–6112 (2011).

22. Andersson, K.M. & Stover, J. The potential impact of a moderately effective HIV vaccine with rapidly waning protection in South Africa and Thailand. Vaccine 29, 6092-6099 (2011).

23. Gray, R.T., Ghaus, M.H., Hoare, A. & Wilson, D.P. Expected epidemiological impact of the introduction of a partially effective HIV vaccine among men who have sex with men in Australia. Vaccine 29, 6125–6129 (2011).

24. Hontelez, J.A., et al. The potential impact of RV144-like vaccines in rural South Africa: a study using the STDSIM microsimulation model. Vaccine 29, 6100–6106 (2011).

25. Long, E.F. & Owens, D.K. The cost-effectiveness of a modestly effective HIV vaccine in the United States. Vaccine 29, 6113–6124 (2011).

26. Nagelkerke, N.J., Hontelez, J.A. & de Vlas, S.J. The potential impact of an HIV vaccine with limited protection on HIV incidence in Thailand: a modeling study. Vaccine 29, 6079–6085 (2011).

27. Phillips, A.N., et al. Potential future impact of a partially effective HIV vaccine in a southern African setting. PLoS One 9, e107214 (2014).

28. Schneider, K., Kerr, C.C., Hoare, A. & Wilson, D.P. Expected epidemiological impacts of introducing an HIV vaccine in Thailand: a model-based analysis. Vaccine 29, 6086-6091 (2011).

29. Darwin, C. On the Origin of Species by Means of Natural Selection, or the Preservation of Favoured Races in the Struggle for Life, (John Murray, London, 1859).

30. Eaton, J.W., et al. HIV treatment as prevention: systematic comparison of mathematical models of the potential impact of antiretroviral therapy on HIV incidence in South Africa. PLoS medicine 9, e1001245 (2012).

31. UNAIDS. Prevention Gap Report. (2016).

32. Cohen, T., Colijn, C. & Murray, M. Modeling the effects of strain diversity and mechanisms of strain competition on the potential performance of new tuberculosis vaccines. Proc Natl Acad Sci U S A 105, 16302–16307 (2008).

33. Pitzer, V.E., et al. Modeling rotavirus strain dynamics in developed countries to understand the potential impact of vaccination on genotype distributions. Proc Natl Acad Sci U S A 108, 19353–19358 (2011).

34. Lipsitch, M. Bacterial vaccines and serotype replacement: lessons from Haemophilus influenzae and prospects for Streptococcus pneumoniae. Emerg Infect Dis 5, 336–345 (1999).

35. Bottomley, C., Roca, A., Hill, P.C., Greenwood, B. & Isham, V. A mathematical model of serotype replacement in pneumococcal carriage following vaccination. J R Soc Interface 10, 20130786 (2013).

36. McLean, A.R. Vaccination, evolution and changes in the efficacy of vaccines: a theoretical framework. Proc Biol Sci 261, 389–393 (1995).

37. Lehman, D.A., et al. Risk of drug resistance among persons acquiring HIV within a randomized clinical trial of single- or dual-agent preexposure prophylaxis. J Infect Dis 211, 1211–1218 (2015).

38. TenoRes Study, G. Global epidemiology of drug resistance after failure of WHO recommended first-line regimens for adult HIV-1 infection: a multicentre retrospective cohort study. The Lancet. Infectious diseases 16, 565–575 (2016).

39. Dimitrov, D., Kublin, J.G., Ramsey, S. & Corey, L. Are Clade Specific HIV Vaccines a Necessity? An Analysis Based on Mathematical Models. EBioMedicine 2, 2062–2069 (2015)

40. Rolland, M., Nickle, D.C. & Mullins, J.I. HIV-1 group M conserved elements vaccine. PLoS Pathog 3, e157 (2007).

41. Rademeyer, C., et al. Features of Recently Transmitted HIV-1 Clade C Viruses that Impact Antibody Recognition: Implications for Active and Passive Immunization. PLoS Pathog 12, e1005742 (2016).

42. Bunnik, E.M., et al. Adaptation of HIV-1 envelope gp120 to humoral immunity at a population level. Nat Med 16, 995–997 (2010).

43. Kawashima, Y., et al. Adaptation of HIV-1 to human leukocyte antigen class I. Nature 458, 641–645 (2009).

44. Carlson, J.M., et al. Impact of pre-adapted HIV transmission. Nat Med 22, 606–613 (2016)

45. Fryer, H.R. & McLean, A.R. Modelling the spread of HIV immune escape mutants in a vaccinated population. PLoS Comput Biol 7, e1002289 (2011).

46. Nowak, M.A. & McLean, A.R. A mathematical model of vaccination against HIV to prevent the development of AIDS. Proc Biol Sci 246, 141–146 (1991).

47. Mullins, J.I., Rolland, M. & Allen, T.M. Viral evolution and escape during primary human immunodeficiency virus-1 infection: implications for vaccine design. Curr Opin HIV AIDS 3, 60–66 (2008).

48. Chen, R.T., et al. Preparing for the availability of a partially effective HIV vaccine: some lessons from other licensed vaccines. Vaccine 29, 6072–6078 (2011).

49. Herbeck, J.T., et al. Evolution of HIV virulence in response to widespread scale up of antiretroviral therapy: a modeling study. Virus Evolution 2, vew028 (2016).

50. Herbeck, J.T., Mittler, J.E., Gottlieb, G.S. & Mullins, J.I. An HIV epidemic model based on viral load dynamics: value in assessing empirical trends in HIV virulence and community viral load. PLoS Comput Biol 10, e1003673 (2014).

51. Jenness, S.M., et al. Impact of the Centers for Disease Control's HIV Preexposure Prophylaxis Guidelines for Men Who Have Sex With Men in the United States. J Infect Dis (2016).

52. Health, S.A.D.o. The 2010 National Antenatal Sentinel HIV and Syphilis Prevalence Survey in South Africa. (Pretoria, South Africa, 2011).

53. Cori, A., et al. CD4(+) cell dynamics in untreated HIV-1 infection: overall rates, and effects of age, viral load, sex and calendar time. Aids 29, 2435–2446 (2015).

54. Fraser, C., Hollingsworth, T.D., Chapman, R., de Wolf, F. & Hanage, W.P. Variation in HIV-1 set-point viral load: Epidemiological analysis and an evolutionary hypothesis. Proceedings of the National Academy of Sciences of the United States of America 104, 17441–17446 (2007).

55. Handcock, M.S., Hunter, D.R., Butts, C.T., Goodreau, S.M. & Morris, M. statnet: Software Tools for the Representation, Visualization, Analysis and Simulation of Network Data. J Stat Softw 24, 1548–7660 (2008).

56. Krivitsky, P.N. Exponential-family random graph models for valued networks. Electron J Stat 6, 1100–1128 (2012).

57. Hughes, J.P., et al. Determinants of Per-Coital-Act HIV-1 Infectivity Among African HIV-1-Serodiscordant Couples. Journal of Infectious Diseases 205, 358–365 (2012).

58. Jenness, S.M., Goodreau, S.M. & Morris, M. EpiModel: Mathematical Modeling of Infectious Disease. (2016).

59. Gilbert, P., Dimitrov, D. & Selinger, C. in 4th Annual Institute for Disease Modeling Symposium (Bellevue, WA, 2016).

60. Pilcher, C.D., et al. Brief but efficient: acute HIV infection and the sexual transmission of HIV. J Infect Dis 189, 1785–1792 (2004).

61. Schacker, T.W., Hughes, J.P., Shea, T., Coombs, R.W. & Corey, L. Biological and virologic characteristics of primary HIV infection. Ann Intern Med 128, 613–620 (1998).

62. Geskus, R.B., et al. The HIV RNA setpoint theory revisited. Retrovirology 4(2007).

63. Herbeck, J.T., et al. Is the virulence of HIV changing? A meta-analysis of trends in prognostic markers of HIV disease progression and transmission. Aids 26, 193–205 (2012).

64. Korenromp, E.L., Williams, B.G., Schmid, G.P. & Dye, C. Clinical Prognostic Value of RNA Viral Load and CD4 Cell Counts during Untreated HIV-1 Infection-A Quantitative Review. Plos One 4 (2009).

65. Fraser, C., et al. Virulence and Pathogenesis of HIV-1 Infection: An Evolutionary Perspective. Science 343, 1328-+ (2014).

66. Little, S.J., McLean, A.R., Spina, C.A., Richman, D.D. & Havlir, D.V. Viral dynamics of acute HIV-1 infection. J Exp Med 190, 841–850 (1999).

67. Fiebig, E.W., et al. Dynamics of HIV viremia and antibody seroconversion in plasma donors: implications for diagnosis and staging of primary HIV infection. Aids 17, 1871-1879 (2003).

68. Piatak, M., et al. High levels of HIV-1 in plasma during all stages of infection determined by competitive PCR. Science 259, 1749–1754 (1993).

69. Patel, P., et al. Estimating per-act HIV transmission risk: a systematic review. Aids 28, 1509–1519 (2014).

70. Goodreau, S.M., et al. What drives the US and Peruvian HIV epidemics in men who have sex with men (MSM)? Plos One 7, e50522 (2012).

71. Palmer, S., et al. New real-time reverse transcriptase-initiated PCR assay with singlecopy sensitivity for human immunodeficiency virus type 1 RNA in plasma. J Clin Microbiol 41, 4531–4536 (2003).

